# Five Breaks Synchrony While Six Keeps Synchrony: Individual Difference in the Coordinated Pattern of Five- and Six-armed Brittle Stars

**DOI:** 10.1101/340471

**Authors:** Daiki Wakita, Yumino Hayase, Hitoshi Aonuma

**Affiliations:** Graduate School of Life Science, Hokkaido University, Sapporo 060-0810, Japan; Graduate School of Science, Hiroshima University, Higashi-Hiroshima 739-8526, Japan; Research Institute for Electronic Science, Hokkaido University, Sapporo 060-0812, Japan

## Abstract

Physiological experiments and mathematical models have supported that neuronal activity is crucial for coordinating rhythmic movements in animals. On the other hand, robotics studies have suggested the importance of physical properties made by body structure, i.e. morphology. However, it remains unclear how morphology affects movement coordination in animals, independent of neuronal activity. To begin to understand this issue, our study reports a rhythmic movement in the green brittle star. We found this animal moved five radially symmetric parts in a well-ordered unsynchronized pattern. We built a phenomenological model where internal fluid flows between the five body parts to explain the coordinated pattern without considering neuronal activity. Changing the number of the body parts from five to six, we simulated a synchronized pattern, which was demonstrated also by an individual with six symmetric parts. Our model suggests a different number in morphology makes a different fluid flow, leading to a different synchronization pattern in the animal.

## Introduction

Animals exhibit a variety of rhythmic movements, such as locomotion, mastication and ventilation. It is generally accepted that animals utilize neuronal activity to coordinate the movements of distant body parts, such as left and right limbs in walking. Many studies have demonstrated the relationship between neuronal interactions and movement coordination by mathematical models. One such successful model to explain the swimming patterns of lampreys was built based on the results of physiological experiments and simulated swimming movements^1^. A genetics study using transgenic mice experimentally changed the coordinated patterns of left and right limbs to reveal the role of specific neurons in locomotion^2^. On the other hand, robotics studies have suggested the importance of physical properties in body for such coordination. For example, four-limbed robots demonstrated different locomotion patterns depending on the loading weight or the swing speed of each motor-driven limb although the limbs were physically connected without electrical circuitry (i.e. neuronal circuitry in animals)^3–4^. A brittle-star-like robot designed by a similar concept was able to immediately change locomotion patterns owing to the unexpected loss of limbs^5^. In these robots, the coordinated patterns of body parts are directly influenced by body structure, i.e. morphology. Supposedly, such a morphological role independent of electricity is also utilized in animals as non-neuronal interactions. However, it remains an issue of how morphology affects such coordinated movements in animals. This problem arises from the lack of examples in which obviously different morphology results in obviously different coordinated patterns in animals.

To break this impasse, we report a rhythmic movement in the green brittle star *Ophiarachna incrassata* (Lamarck, 1816). This marine animal is characterized by radial symmetry in morphology, as recognized in other echinoderms. The central disk has typically five arms, which partition the disk into five symmetric fan-shaped parts, called interradii. The movement we found is the rhythmic shrinkage and expansion of the five interradii and frequently occurs after feeding (Fig. 1, Movie S1). Individuals that were kept without feeding even for a week never showed the movement, implying a relationship with digestion. It was reminiscent of pumps, hence we termed this rhythmic movement “*pumping*”. The movements between the five interradii were regularly ordered without synchronization. Based on our mathematical model, this coordinated pattern can be easily explained by physical properties without considering neuronal interactions. We then changed the number of interradii in the model, referring to the fact that brittle stars sometimes have individual difference in the number of arms, namely, the number of interradii. Changing from five to six interradii simulated synchronized movements between the interradii (Fig 2). We finally obtained a six-armed individual, which demonstrated the synchronized movements (Fig. 3, Movie S2). Therefore pumping will provide a model system to understand the role of morphology in movement coordination, utilized in actual animals.

**Fig. 1.**
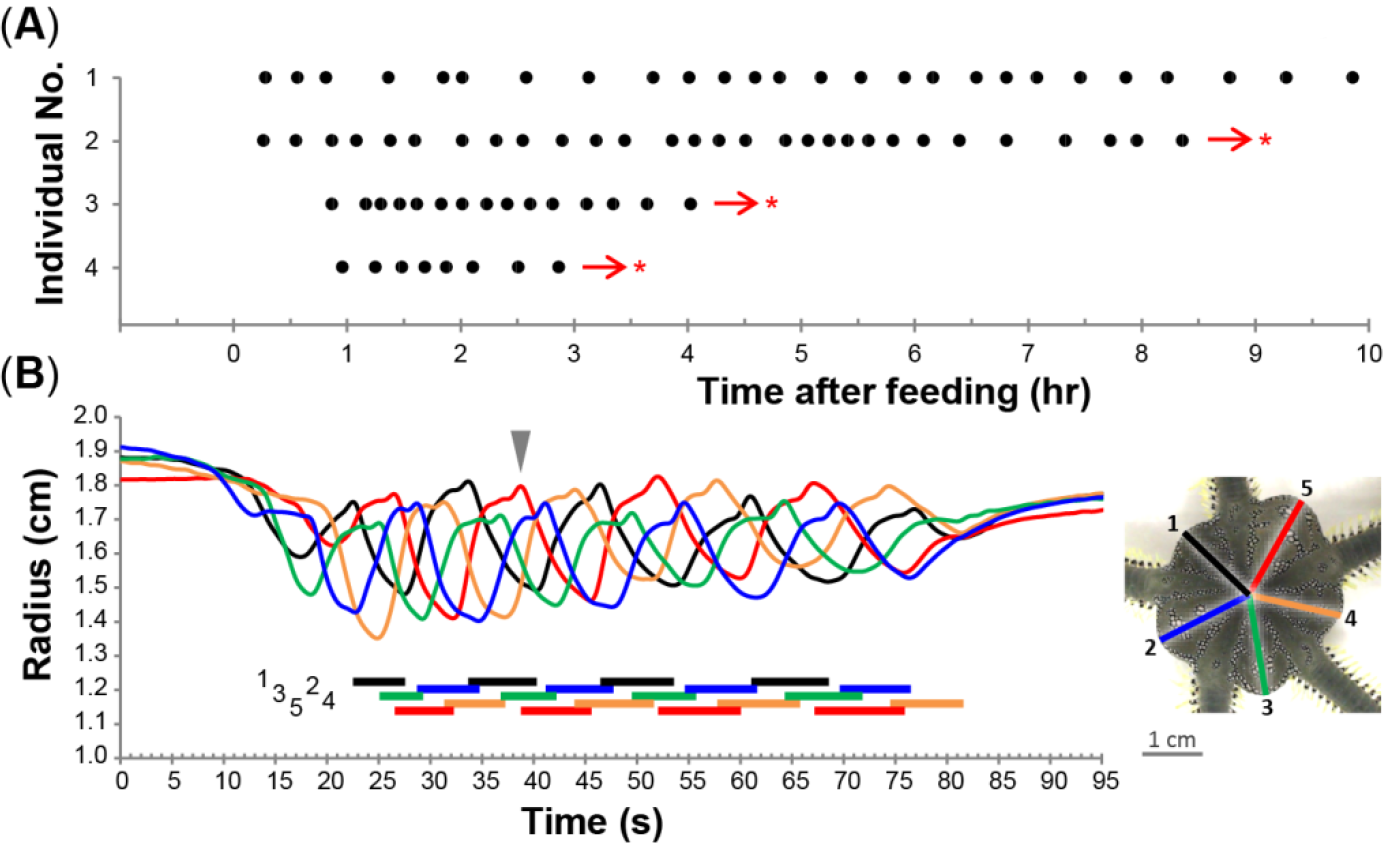
Rhythmic movement, “pumping”, in the five-armed individuals of the green brittle star *Ophiarachna incrassata*. (**A**) Temporal frequency of pumping phases. Each point represents a pumping phase, which comprises a series of movements shown in (B). The animals exhibit the first pumping phase 36 ± 17 min (N = 4) after feeding. Then, they periodically initiate pumping for more than 10 hrs. The interval between pumping phases is not consistent (20 ± 9 min) among individuals. Asterisks indicate no record from the arrows. (**B**) Temporal change in the radius of the five interradii in a pumping phase. Inset shows the aboral side of an individual at the moment indicated by the arrowhead in the graph. Radius is measured from the center of the disk to the midpoint of the edge of each interradius. The radii numbered anticlockwise are colored as in the inset, which corresponds to the graph in color. Colored horizontal bars under the graph represent shrinking periods of each interradius.

**Fig. 2.**
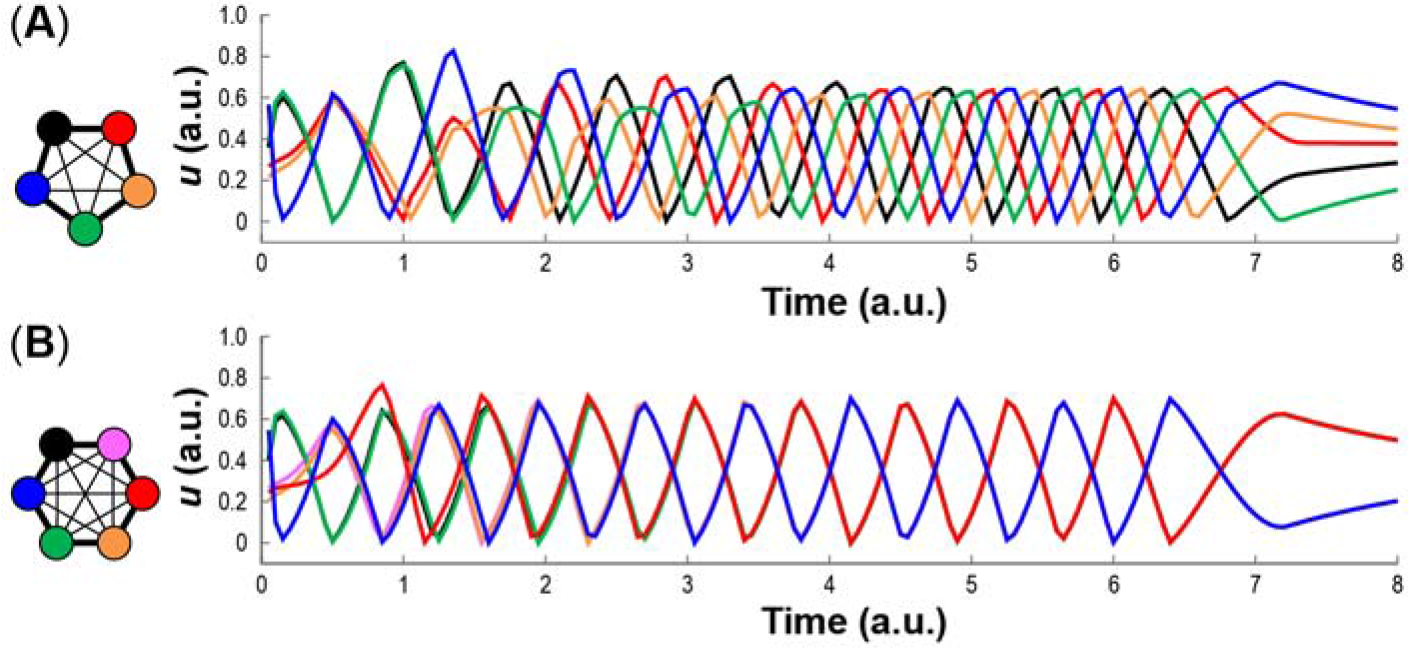
Simulation of rhythmic movement, “pumping” in the green brittle star *Ophiarachna incrassata*, based on a phenomenological model. (**A**) Temporal change of the volume of five interradii. Cycles are unsynchronized as in the experiments of five-armed individuals (see Fig. 1B). (**B**) Temporal change of the volume of six interradii. Three distant interradii and the other three each make synchronized groups, anticipating the coordinated pattern of six-armed individuals. *u* in the y-axis represents the volume of internal fluid in the interradii. Each axis is given in an arbitrary unit. The color of the interradii in each inset corresponds to that of each graph.

**Fig. 3.**
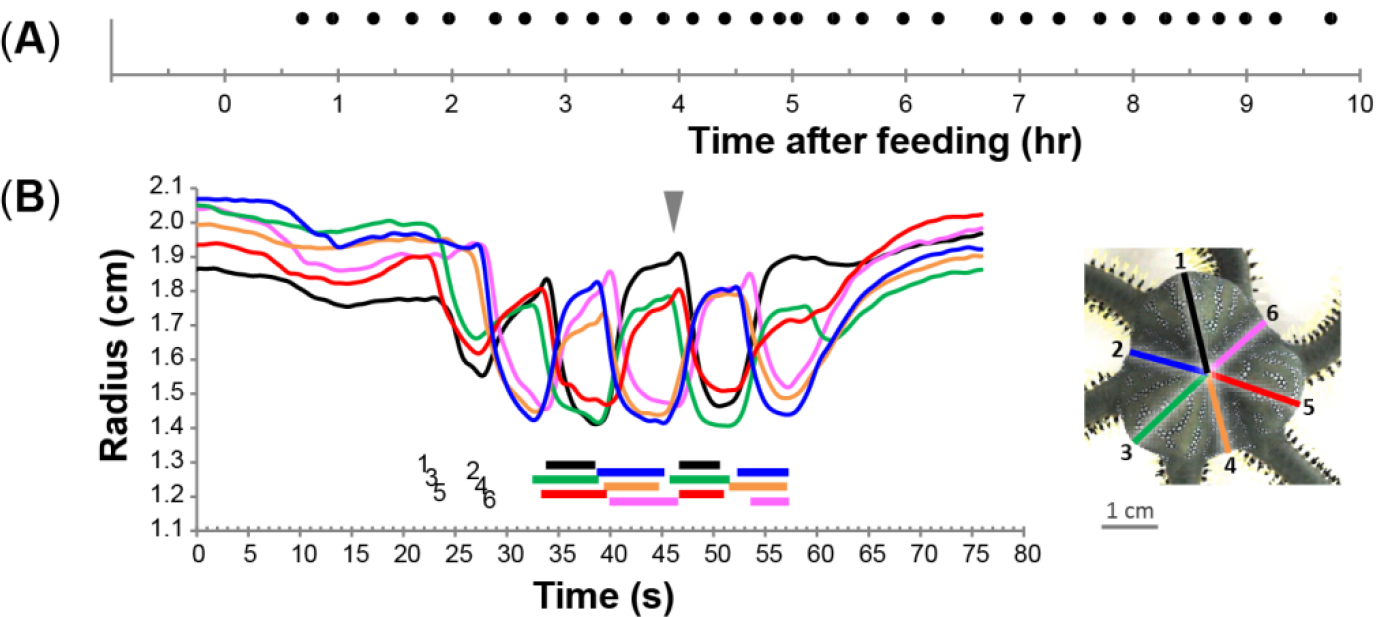
Rhythmic movement, “pumping”, in the six-armed individual of the green brittle star *Ophiarachna incrassata*. (**A**) Temporal frequency of pumping phases. The animal frequently shows pumping phases for more than 10 hrs after feeding, as occurs in five-armed animals. (**B**) Temporal change in the radius of the six interradii in a pumping phase. Cycles are synchronized with two groups separated, which demonstrates the model simulation of six interradii (see Fig. 2B). Figures are shown as in Fig. 1.

## Results and Discussion

We here describe the rhythmic movement, pumping, in the green brittle star. Pumping was frequently observed with an interval of 21 ± 10 min (mean ± S.D., N = 4) after feeding (Fig. 1A). Each continuously moving phase, termed “*pumping phase*”, started with a closing of the mouth and genital slits on the oral side of the disk. The whole contraction of the disk was immediately followed by a series of shrinkage and expansion in interradii. Five-armed individuals produced a pumping phase for 57 ± 12 s (N = 2, n = 6), in which each of the five interradii repeated 3–5 cycles of the movements. Based on the radius of the interradii in the aboral view, each cycle comprised a shrinking period for 5.9 ± 1.0 s and an expanding period for 6.5 ± 0.8 s at the beginning of the pumping phase (Fig. 1B). These phases gradually became longer to 7.6 ± 1.0 and 8.1 ± 0.9 s respectively at the end. Meanwhile, the amplitude of the movements gradually became smaller. Each expanding period included two increasing stages in radius; the first longer one was recognized as an increase in the volume of the interradii, while the second shorter one was apparently ascribed to change in the form of the interradii. Numbered from 1 to 5 anticlockwise in the aboral view, the interradii regularly moved in the unsynchronized sequence of 1-3-5-2-4 or 1-4-2-5-3 repetitively; a cycle in one of the five interradii was followed by that in either of two distant ones with 2.2 ± 0.5 s delayed (Fig. 1B, Movie S1). The two types of sequence were observed in the same individual although a continuous pumping phase underwent either one consistently. The initially shrinking interradius was not always identical. At the end of the pumping phase, the mouth and genital slits slowly opened, so that the disk became relaxed as usual.

To explain the relationship between the pumping pattern and morphology, we constructed a mathematical model. Previous models explained how differences in network structure caused difference in rhythmic outputs on the basis of neuronal networks where a certain number of neurons constituted a ring^6–8^. These models were inspired in contexts irrelevant to pumping; nevertheless, they demonstrated that a pumping-like rhythmic pattern was generated among five neurons which laterally inhibited each other. Although brittle stars possess a nerve ring in the disk^9^, it is unclear whether they utilize such a neuronal network. We suppose that their simple nervous system is insufficient to coordinate the dynamic movement of pumping, and rather physical properties of their body have a dominant effect. We thus construct a phenomenological model simply focusing on the morphological change in interradii, independent of neuronal interactions.

We assume that internal fluid is saved in each interradius with the volume of *u*_*i*_ (*t*) individually. *i* takes 1 to N anticlockwise; *N* = 5 in the case of five interradii. During the shrinking period, the tension of the surface skin increases and the fluid is pushed out to another interradius. We define the pressure *P*_*i*_ (*t*) at each interradius and the fluid flow from higher pressure to lower pressure.

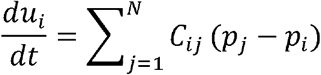

The total volume of fluid 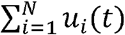 is constant in time. *C*_*ij*_is the constant indicating the connection between i-th and j-th interradii. We assume here, the connection between nearest neighbors is larger than the others.

During the expanding period, the fluid flows into the interradius. We suppose that the volume of fluid increases the pressure similarly to an elastic membrane,

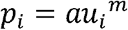

For simplicity *m* 1. During the shrinking period, the interradius contracts to push the fluid out. We suppose a target pressure *p̄*, which is much larger than that of the expanding period.

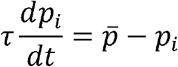

The expanding period switches to the shrinking period because of a capacity limit of *u*_*i*_ (*t*). The switching occurs when *p*_*i*_ (*t*) exceeds the threshold *p*_*th*_, (*p*_*th*_ < *p̄*). On the other hand, the shrinking period switches to the expanding one when no fluid remains, i.e. *u*_*i*_ (*t*) = 0.

At the beginning of the pumping phase, an interradius starts to shrink casually. We assume that a food exists in one of the interradii. We take the initial condition that an interradius has a higher pressure and more fluid than the others: (*u*_1_, *p*_1_) = (*u*_*h*_, *p*_*h*_) and (*u*_*i*_, *p*_*i*_) = (*u*_0_ + *ξ*_*i*_, 0) for *i* = 2, *N*. Here, *u*_*o*_ < *u*_*h*_, *p*_*h*_ ~*p̄* and *ξ*_*i*_ is noise. At the end of the pumping phase, we suppose that the target pressure *p̄* becomes smaller, given the observation that the disk becomes relaxed. With suitably chosen parameters, our numerical simulations support the experimental results well (Fig. 2A).

We focus on the fact that brittle stars sometimes have individual difference in the number of arms, namely, the number of interradii. Although five-armed animals are most common, there are six-armed animals in some cases. Simply changing the number of interradii from five to six leads to a different synchronization pattern in the model simulation (Fig. 2B). Numbered from 1 to 6, the interradii moved in the synchronized sequence of 1, 3 and 5 together and then 2, 4 and 6 together repetitively; synchronized cycles in three distant ones followed that in the other three.

The cause of this difference can be explained by the volume and the pressure. For the case of *N* = 5, the 1st interradius pushes most fluid into the 2nd and 5th ones, and then the 2nd and 5th push it into the 3rd and 4th, respectively. At the beginning, the difference between *u*_3_ and *u*_4_ is small. However, for example, when *p*_3_ exceeds the threshold *p*_th_ and *p*_4_ is still in the expanding period, the fluid flows from the 3rd to the 4th. The difference between *u*_3_ and *u*_4_ increases rapidly (around t = 1.5 in Fig. 2), so that each interradius in five-armed individuals makes an asymmetric flow into either neighbor. The direction of the rotation depends on the initial perturbation. For the case of *N* = 6, the fluid flows started from the 1st interradius are propagated anticlockwise (1-2-3-4) and clockwise (1-6-5-4) and collide with each other at the 4th one. After the collision, the 4th pushes the fluid back to the 3rd and 5th. If the 3rd has a smaller volume than the 5th, the pressure of the 3rd is smaller than that of the 5th because both are in the expanding period. More fluid flows into the 3rd than into the 5th. As a result, *u*_3_ and *u*_5_ come to synchronize. The 4th acts as a buffer to eliminate the difference between *u*_3_ and *u*_5_. This mechanism causes each interradius in six-armed individuals to make symmetric flows into both neighbors, dividing the interradii into two synchronized groups.

After *t* = 6, we decrease the target pressure *p̄* gradually to zero. In both the case of *N* = 5 and 6, the cycles become longer and finally the oscillation vanishes in the same way as in the experiments (Fig. 2).

We then obtained a six-armed individual with six symmetric interradii, in which pumping was also recognized. The interval between pumping phases was 19 ± 2 min (N = 1) after feeding (Fig. 3A). Each of the pumping phases continued for 39 ± 14 s (N = 1, n = 3) with the six interradii each repeating 2–4 cycles. The duration of shrinking and expanding periods at the beginning of the pumping phase was 5.4 ± 0.8 and 6.7 ± 0.6 s respectively, which became 5.7 ± 0.8 and 8.6 ±1.6 s, respectively, at the end. The movements in the interradii were synchronized as occurs in the simulation (Fig. 3B, Movie S2).

Suppose pumping functions in a digestive process, the difference in coordinated pattern can be referred to intestinal movements. The interradii of five-armed individuals can make a travelling wave, which resembles peristaltic motion for transporting liquid matter in intestine^10^. On the other hand, the interradii of six-armed individuals can bring a stationary wave as recognized in segmental motion for mixing solid matter^10^. Accordingly, this individual difference in the brittle star can cause even a functional difference.

The correspondence between the simulation and the real phenomena in two patterns suggests the modelled non-neuronal interaction is likely to coordinate the movements in the animal. Previous studies explaining rhythmic movements in animals have built mathematical models assuming the existence of autonomous oscillatory activity of pacemaker neurons, so-called central pattern generators (CPGs)^1, 11–14^. Robotics studies focusing on physical properties have also employed the idea of CPGs, as designed in each limb for locomotion^3, 4^ and in intestinal wall for peristalsis^15^. Even apart from animals, protoplasmic movement in the true slime mold has inspired physicists and robotics researchers to build mathematical models related to the number of circularly arranged oscillators^16–17^. Although this unicellular organism has no nervous system, these models have instead assumed spatially distributed biochemical oscillators in the body, which are similar to CPGs in concept. In this perspective, our model without explicit CPGs helps us begin to understand how biological movements take advantage of morphology, independent of spontaneous oscillation in neurons or biochemicals.

## Methods

Intact adult individuals of the green brittle star *Ophiarachna incrassata* (Lamarck, 1816) were used in this study. They were obtained commercially and reared in laboratory aquariums (600 mm × 600 mm × 600 mm) filled with artificial seawater at 25–28°C (TetraMarin Salt Pro, Tetra Japan Co, Tokyo, Japan; salinity, 32–35‰). The disks ranged 2–5 cm in diameter and mostly had five symmetric arms. We also obtained an individual that had six arms with a 4 cm diameter disk.

All animals were fed with dried krill (Tetra Krill-E, Tetra Japan Co, Tokyo, Japan). Once they detected the food, their arms captured it and carried it to their mouths. After a while, their disks start moving rhythmically. The long-term behavior of four randomly selected five-armed individuals and one six-armed animal were recorded using a time-lapse camera (TCL200, Brinno, Taipei, Taiwan). The short-term rhythmic movement of two randomly selected five-armed animals and the one six-armed animal were recorded three times for each from the aboral side with a video camera (EOS8000D, Canon, Tokyo, Japan) in small acrylic cases. Successfully recorded movements were analysed using a video tracking software Kinovea (v. 0.8.15, http://www.kinovea.org/) at 10 f.p.s.

The numerical simulations of the mathematical model were carried out for *a* = 1, *p̄* = 20, *p*_*th*_ = 0.5, τ = 1, *C*_*ij*_ = 0.2 for nearest neighbors, *C*_*ij*_ = 0.02 for the others. For the initial condition, *u*_*o*_ = 0.2, *u*_*h*_ = 1, *p*_*h*_ = *p̄*. After *t* = 6, the value of *p̄* is gradually reduced to zero.

## Data Availability

The datasets generated during and/or analyzed during the current study are available from the corresponding author on reasonable request.

## Acknowledgments

We thank Prof. Philip L. Newland for his critical comment upon our manuscript. This work was partly supported by JSPS KAKENHI (Grant Number 16KT0099) and by JST CREST (Grant Number JPMJCR14D5), Japan.

## Author Contributions

D.W., Y.H. and H.A. observed and analyzed the rhythmic movement. D.W., Y.H. and H.A. built the mathematical model. D.W., Y.H. and H.A. wrote the manuscript.

## Supplementary Information

**Movie S1.** Video of pumping in a five-armed individual in the aboral side.

**Movie S2.** Video of pumping in a six-armed individual in the aboral side.

## References

1. Grillner, S. Biological pattern generation: the cellular and computational logic of networks in motion. Neuron 52, 751–766 (2006).

2. Talpalar, A. E. et al. Dual-mode operation of neuronal networks involved in left-right alternation. Nature 500, 85 (2013).

3. Owaki, D., Kano, T., Nagasawa, K., Tero, A. & Ishiguro, A. Simple robot suggests physical interlimb communication is essential for quadruped walking. J. R. Soc. Interface 10, 20120669 (2013).

4. Owaki, D. & Ishiguro, A. A quadruped robot exhibiting spontaneous gait transitions from walking to trotting to galloping. Sci. Rep.-UK 7, 277 (2017).

5. Kano, T., et al. A brittle star-like robot capable of immediately adapting to unexpected physical damage. Roy. Soc. Open Sci. 4, 171200 (2017).

6. Kling, U. & Székely, G. Simulation of rhythmic nervous activities. Kybernetik 5, 89–103 (1968).

7. Suzuki, R., Katsuno, I. & Matano, K. Dynamics of “neuron ring”. Kybernetik 8, 39–45 (1971).

8. Matsuoka, K. Sustained oscillations generated by mutually inhibiting neurons with adaptation. Biol. Cybern. 52, 367–376 (1985).

9. Cobb, J. L. S. & Stubbs, T. R. The giant neurone system in Ophiuroids III: the detailed connections of the circumoral nerve ring. Cell Tissue Res. 226, 675–687 (1982).

10. Umetani, Y. & Inou, N. Biomechanical study of peristalsis: neural mechanism of rhythmic segmentation. J. Soc. Inst. Contr. Eng. 21, 965–969 (1985). [in Japanese]

11. Stein, P. S. Motor systems, with specific reference to the control of locomotion. Ann. Rev. Neurosci. 1, 61–81 (1978).

12. Delcomyn, F. Neural basis of rhythmic behavior in animals. Science 210, 492–498 (1980).

13. Grillner, S., McClellan, A. & Perret, C. Entrainment of the spinal pattern generators for swimming by mechano-sensitive elements in the lamprey spinal cord in vitro. Brain Res. 217, 380–386 (1981).

14. Marder, E. & Calabrese, R. L. Principles of rhythmic motor pattern generation. Physiol. Rev. 76, 687–717 (1996).

15. Kano, T., Kawakatsu, T. & Ishiguro, A. Generating situation-dependent behavior: decentralized control of multi-functional intestine-like robot that can transport and mix contents. J. Robot. Mech. 25, 871–876 (2013).

16. Takamatsu, A. et al. Spatiotemporal symmetry in rings of coupled biological oscillators of Physarum plasmodial slime mold. Phys. Rev. Lett. 87, 078102 (2001).

17. Umedachi, T., Idei, R., Ito, K. & Ishiguro, A. A fluid-filled soft robot that exhibits spontaneous switching among versatile spatiotemporal oscillatory patterns inspired by the true slime mold. Artif. Life 19, 67–78 (2013).

